# Impact and evolutionary determinants of Neanderthal introgression on transcriptional and post-transcriptional regulation

**DOI:** 10.1101/532366

**Authors:** Martin Silvert, Lluis Quintana-Murci, Maxime Rotival

## Abstract

Archaic admixture is increasingly recognized as an important source of diversity in modern humans, with Neanderthal haplotypes covering 1-3% of the genome of present-day Eurasians. Recent work has shown that archaic introgression has contributed to human phenotypic diversity, mostly through the regulation of gene expression. Yet, the mechanisms through which archaic variants alter gene expression, and the forces driving the introgression landscape at regulatory regions remain elusive. Here, we explored the impact of archaic introgression on transcriptional and post-transcriptional regulation, focusing on promoters and enhancers across 127 different tissues as well as microRNA-mediated regulation. Although miRNAs themselves harbor few archaic variants, we found that some of these variants may have a strong impact on miRNA-mediated gene regulation. Enhancers were by far the regulatory elements most affected by archaic introgression, with one third of the tissues tested presenting significant enrichments. Specifically, we found strong enrichments of archaic variants in adipose-related tissues and primary T cells, even after accounting for various genomic and evolutionary confounders such as recombination rate and background selection. Interestingly, we identified signatures of adaptive introgression at enhancers of some key regulators of adipogenesis, raising the interesting hypothesis of a possible adaptation of early Eurasians to colder climates. Collectively, this study sheds new light onto the mechanisms through which archaic admixture have impacted gene regulation in Eurasians and, more generally, increases our understanding of the contribution of Neanderthals to the regulation of acquired immunity and adipose homeostasis in modern humans.

## Introduction

The sequencing of the genomes of extinct human forms, such as Neanderthals or Denisovans, has enabled the mapping of archaic variants in the genomes of modern humans.^1–7^ This archaic introgression has functional consequences today, as introgressed variants have been reported to alter a variety of phenotypes, ranging from skin pigmentation to sleeping patterns and mood disorders.^8,9^ Importantly, there is accumulating evidence to suggest that the impact of Neanderthal introgression on complex, ultimate phenotypes is largely mediated by common variants regulating gene expression.^10,11^ For example, up to a quarter of Neanderthal-introgressed haplotypes have been estimated to present *cis*- regulatory effects across tissues, with a bias towards down-regulation of Neanderthal alleles in brain and testes.^12^ Furthermore, genes involved in innate immunity and interactions with RNA viruses have been reported to be enriched in Neanderthal ancestry,^13,14^ with archaic variants affecting, in particular, transcriptional responses to viral challenges.^11,15^ However, the regulatory mechanisms mediating the effects of Neanderthal variants on gene expression and immune responses remain poorly known.

Our understanding of the selective forces that shaped the landscape of archaic introgression is also rapidly growing. In most cases, archaic variants were selected against, with regions of higher selective constraint being depleted in archaic ancestry, particular those that are X-linked or contain testis-expressed and meiotic-related genes.^1,2,16^ Some studies have also suggested that Neanderthals had a reduced effective population size,^6,7^ owing to a prolonged bottleneck or a deeply structured population.^6,17,18^ Natural selection in Neanderthals would thus have been less efficient to purge deleterious mutations,^19,20^ a large proportion of which were removed from the genome of modern humans following their admixture with Neanderthals.^21^ However, archaic variants have also contributed, in some cases, to human adaptation,^15,22-26^ shortly after their introduction into modern humans or after an initial period of neutral evolution.^27,28^ Given the rapid evolution of regulatory regions and their potential adaptive nature,^29,30^ the evolutionary dynamics of Neanderthal introgression at regulatory elements needs to be explored in further detail.

## Results

In this study, we aimed to increase knowledge on the impact that archaic introgression has had on transcriptional and post-transcriptional mechanisms, focusing on promoter-, enhancer- and microRNA-mediated regulation.^31,32^ To this end, we first characterized the set of variants of putative Neanderthal origin — archaic SNPs (aSNPs) — as those for which one allele is both present in the Neanderthal Altai genome^6^ and absent in the Yoruba African population of the 1000 Genomes project (**Supplemental Methods**).^33^ We further required aSNPs to be located in genomic regions where Neanderthal introgression has already been detected in Europe or Asia.^1^ We then investigated deviations in the presence or absence of aSNPs among specific classes of functional elements. To do so, we measured the density of aSNPs, with respect to that of non-aSNPs, in the European (CEU) and Asian (CHB) populations of the 1000 Genomes project.^33^ We then compared the relative density of aSNPs at specific regulatory regions to that of the rest of the genome. The studied elements were considered as enriched/depleted in aSNPs if the resulting odds ratio was significantly different from 1.

Overall, we observed a strong depletion of aSNPs in coding regions (OR = 0.71, *p*-value<10^-4^), as expected,^1,2,4^ and levels of introgression at regulatory regions were similar to those of non-functional elements. However, when aSNPs were classified according to their minor allele frequency (MAF) — to distinguish rare (MAF < 1%, 21% of aSNPs), low-frequency (1% ≤ MAF < 5%, 48% of aSNPs) and common (MAF ≥ 5%, 31% of aSNPs) variants — the depletion in coding regions was found to be driven by rare and low-frequency variants (OR < 0.79, *p*-value < 2 × 10^-5^). Conversely, regulatory regions were significantly enriched in low-frequency and common aSNPs (OR > 1.04, *p*-value < 0.05; **Figure 1A**), consistent with the notion that aSNPs tend to have regulatory effects.^10,11^ While the relative density of aSNPs varied markedly across frequency bins, it did not differ across promoters, enhancers, and miRNA binding sites, despite their important differences in strength of negative or background selection (**Figures 1B-D**). Neanderthal variants are, however, quantitatively more likely to affect gene regulation via modification of enhancer activity, due to the larger size of enhancers with respect to other regulatory elements (**Figure 1E**).

**Figure 1.**
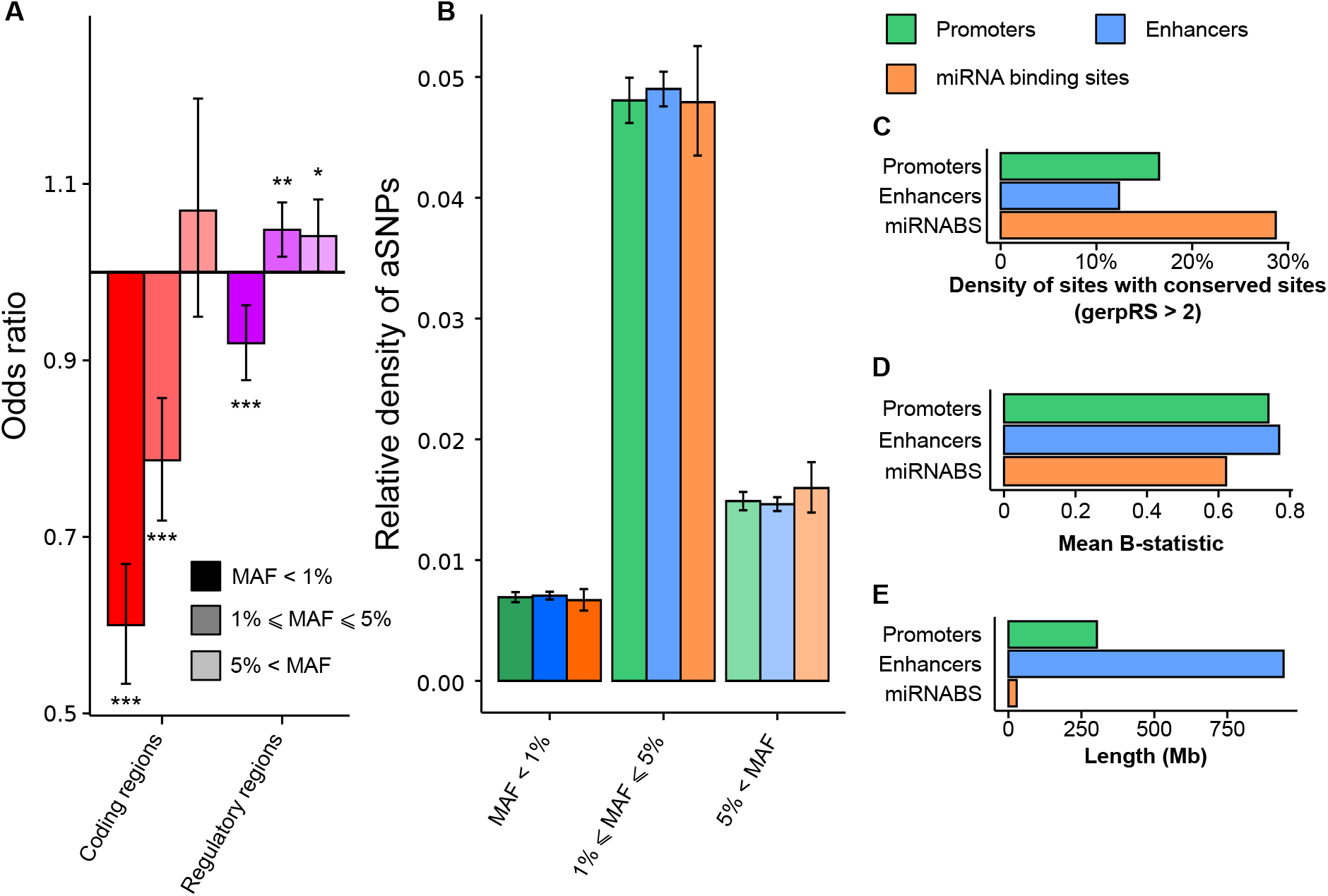
Enrichment of Neanderthal variants in regulatory regions. (A) Odds ratio depicting the excess or depletion of Neanderthal variants in coding regions and regulatory elements (promoters, enhancers and miRNA binding sites), compared to the remainder of the genome. Enrichments are shown for 3 bins of frequencies, with 95% bootstrap confidence intervals: *: *p*-value < 0.05, **: *p*-value <0.01, ***: *p*-value < 0.001. (B) Relative density of aSNPs in promoters, enhancers and miRNA binding sites in different bins of Minor Allele Frequency (MAF), with 95% bootstrap confidence intervals. (C-E) Comparison of density of conserved sites (GerpRS > 2), mean B-statistic and total length of promoters, enhancers, and miRNA binding sites.

We first focused on miRNAs and miRNA binding sites (miRNABS); given the low fraction of the genome that is covered by these elements (**Figure 1E**), they are expected to be, quantitatively, the least affected by archaic introgression. Indeed, we only found 6 aSNPs that overlap the sequence of mature miRNAs, two of which alter the seed region (**Figure 2A**): rs74904371 in miR-2682-3p (MAF_CHB_ = 0, MAF_CEU_ = 3%) and rs12220909 in miR-4293 (MAF_CHB_ = 17%, MAF_CEU_ = 0). The presence of aSNPs in four of these miRNAs, particularly those located in seed regions, affected the set of genes they bind (**Figure 2B and Table S1**). We also detected 2,909 aSNPs in miRNABS, 29% of which were common (**Table S2**). We found a direct linear relationship between the number of genes bound by a miRNA and the number of aSNPs in its binding sites (R^2^ = 31%, *p*-value < 10^-10^; **Figure 2C**), suggesting that introgression affected miRNABS independently of their cognate miRNAs. Highlighting a pertinent example, the *ONECUT2* locus (MIM: 604894) presents the highest number of aSNPs altering conserved miRNABS (**Figure 2D**), and has been previously reported as a likely target of adaptive introgression.^22^ This gene, which encodes a member of the onecut family of transcription factors, contains 13 aSNPs that alter miRNABS, 6 of which are highly conserved (GerpRS > 2). Interestingly, these aSNPs fall within the 0.4% most differentiated aSNPs between Europeans and Asians at the genome-wide level (*F*_ST_ > 0.38). We also detected aSNPs, mostly population specific, that alter conserved miRNABS at several, key immune genes, including *CXCR5* (MIM: 601613; MAF_CHB_ = 16%, MAF_CEU_ = 1%), *TLR6* (MIM: 605403; MAF_CHB_ = 8%, MAF_CEU_ = 0), *IL7R* (MIM: 146661; MAF_CHB_ = 8%, MAF_CEU_ = 0) or *IL21* (MIM: 605384; MAF_CHB_ = 0, MAF_CEU_ = 8%).

**Figure 2.**
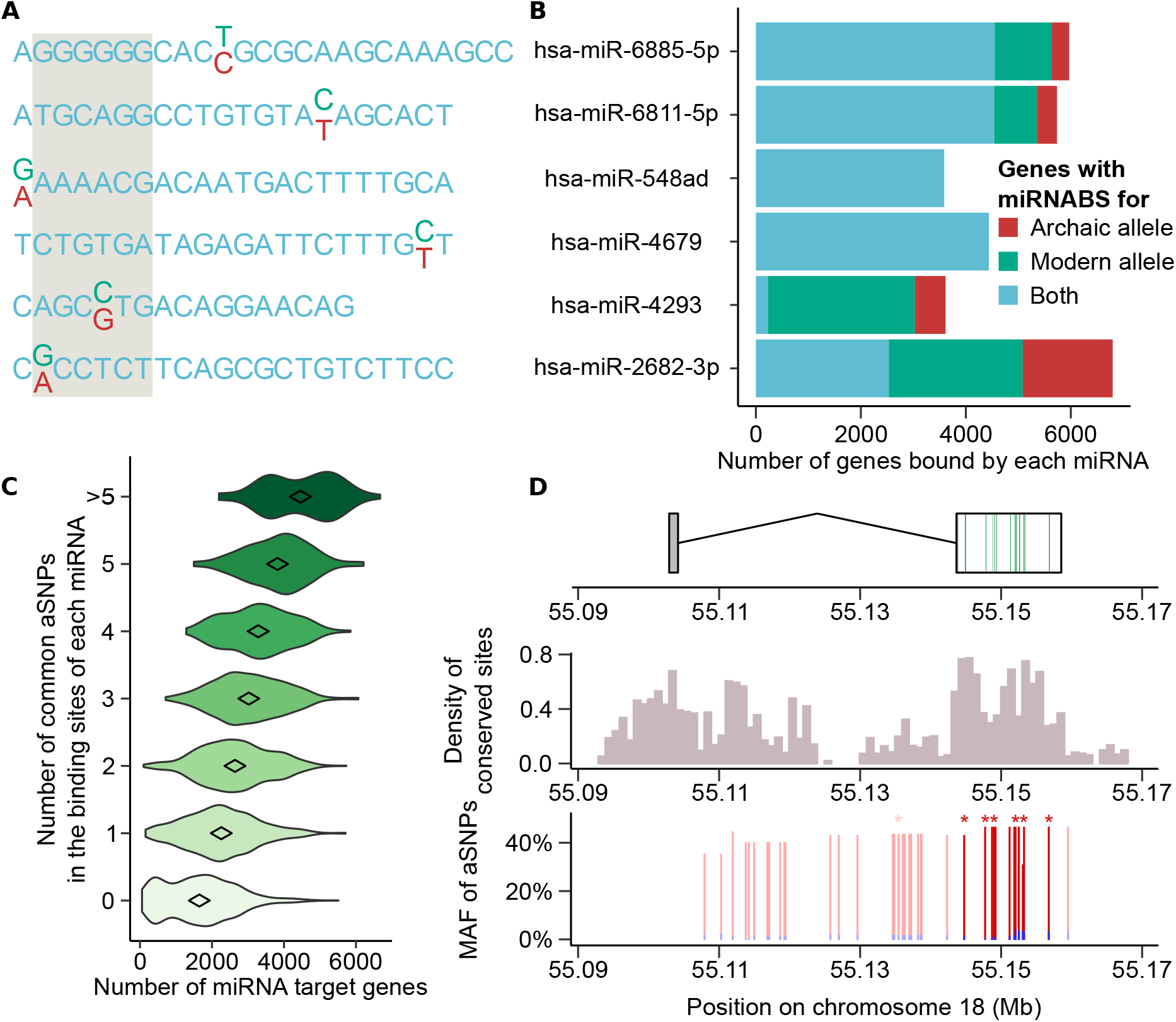
Effects of archaic introgression on miRNA-mediated regulation. (A) Representation of the archaic (red) and modern (green) human alleles for the 6 miRNAs presenting a Neanderthal-introgressed variant in their mature sequence. The seed region of the miRNAs is shaded in grey. (B) Total number of genes bound by the archaic/modern human allele of each of the 6 miRNAs harboring a Neanderthal variant in their mature sequence. (C) Relationship between the number of targets of each miRNA and the number of common aSNPs in the corresponding miRNA binding sites. (D) Introgression of aSNPs altering the miRNABS at the *ONECUT2* locus (MIM: 604894). Gene structure is shown in upper panel, with miRNA binding sites that are altered by archaic introgression highlighted in green. The middle panel represents the density of conserved sites (GerpRS > 2) in 1000bp-windows and the bottom panel the repartition and frequency of archaic alleles at the locus (blue for CEU, red for CHB). aSNPs that overlap miRNABS are represented with a darker shade and aSNPs that disrupt a conserved site are marked with stars.

Next, we focused on how archaic introgression has affected promoters and enhancers. Given the tissue-specific impact of archaic introgression on gene regulation,^10,12^ we searched for enrichments in Neanderthal ancestry across regulatory elements in 127 different tissues.^31^ The impact of archaic introgression in promoters was similar to that of the remainder of the genome, in all tissues and frequency bins (**Table S3**). Conversely, we found that enhancers are enriched in common aSNPs in 42 tissues (FDR < 5%, **Figure 3A and Table S4**). Of these, enhancers that are active in adipose-derived mesenchymal stem cells (AdMSC) and mesenchymal stem cell-derived adipocytes were the most significantly enriched (OR > 1.13, *p*-value < 3×10^-5^), followed by those active in fetal heart (OR = 1.15, *p*-value = 8×10^-5^), small intestine (OR = 1.21, *p*-value = 2×10^-4^) and different T cell tissues (OR > 1.14, *p*-value < 1.5×10^-2^). Focusing on circulating immune cell types (**Figure 3B**), we found enrichments among enhancers of various types of primary T cells, the most significant being CD4+/CD25^-^ memory T cells (OR = 1.21, *p*-value = 2.2×10^-4^), while enhancers of B cells, monocytes and natural killer cells exhibited a density of common aSNPs similar to genome-wide expectations. We also observed that shared enhancers across different T cell subtypes (i.e., active in more than half of T cells subtypes, “core T cells enhancers”) display an enrichment in aSNPs (OR = 1.22, *p*-value = 5×10^-4^, **Figure 3C**), with respect to more specialized enhancers that are only active in a small fraction of T cell subtypes.

**Figure 3.**
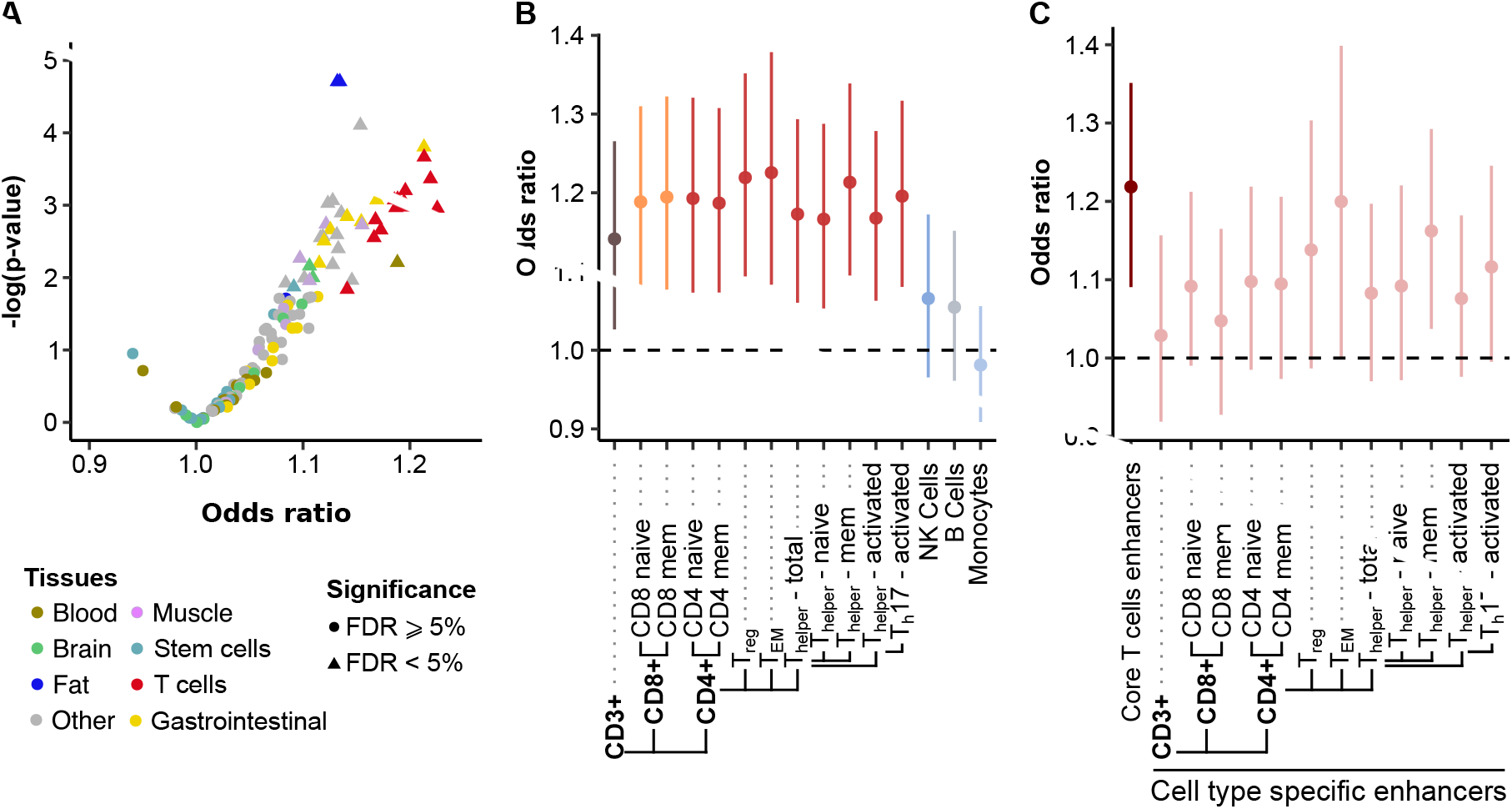
Effects of archaic introgression at enhancers. (A) Volcano plot illustrating the enrichment of common aSNPs in the enhancers of 127different tissues from the Epigenomic Roadmap Consortium. Tissues with an FDR < 5% (triangles) are significantly enriched. (B) Enrichments of common aSNPs in the enhancers of different immune tissues. Vertical bars indicate 95% confidence intervals computed by bootstrap. (C) Enrichment of common aSNPs in the enhancers that are active in more than half of the investigated T cell types (dark red, referred to as “core T cells”) and in enhancers that are active in each T cell type and are not part of core T cell enhancers (light red, referred to as “cell type specific enhancers”). (B,C) Note that CD4^+^ T cells are separated based on CD25 to distinguish Treg (CD25^+^), T_EM_ (CD25^Iow^) and T_helper_ (CD25^-^).

We sought to assess whether the enrichment in aSNPs detected in enhancers resulted from an excessive divergence of these elements in the Neanderthal lineage, or a higher rate of archaic introgression at enhancers. We quantified the number of fixed human/Neanderthal differences at enhancers, across the 127 tissues, focusing on sites where Neanderthals differed from the ancestral sequence. We uncovered a large tissue variability, with enhancers active in induced pluripotent stem cells presenting the highest divergence (290 differences/Mb) and those active in pancreas showing the lowest (220 differences/Mb). However, given that the number of fixed differences strongly correlates with genetic diversity (i.e., density of common variants, r = 0.71, *p*-value <10^-20^), we measured the ratio of the number of fixed human/Neanderthal differences to that of common, segregating SNPs in the region. Using this metric, we found that enhancers of T cells displayed the strongest divergence (7% increase compared to the mean across tissues, Wilcoxon *p*-value < 2×10^-8^), whereas stem cells showed the lowest (4% decrease, Wilcoxon *p*-value < 7×10^-6^) (**Figure 4A**). Focusing on the rate of introgression, defined as the proportion of Neanderthal-descent alleles that are present in the human genome at a MAF > 5%, we found that enhancers of T cells showed the highest percentage (5% increase, Wilcoxon *p*-value < 2×10^-5^), while brain the lowest (7% decrease, Wilcoxon *p*-value < 4×10^-5^) (**Figure 4A**).

**Figure 4.**
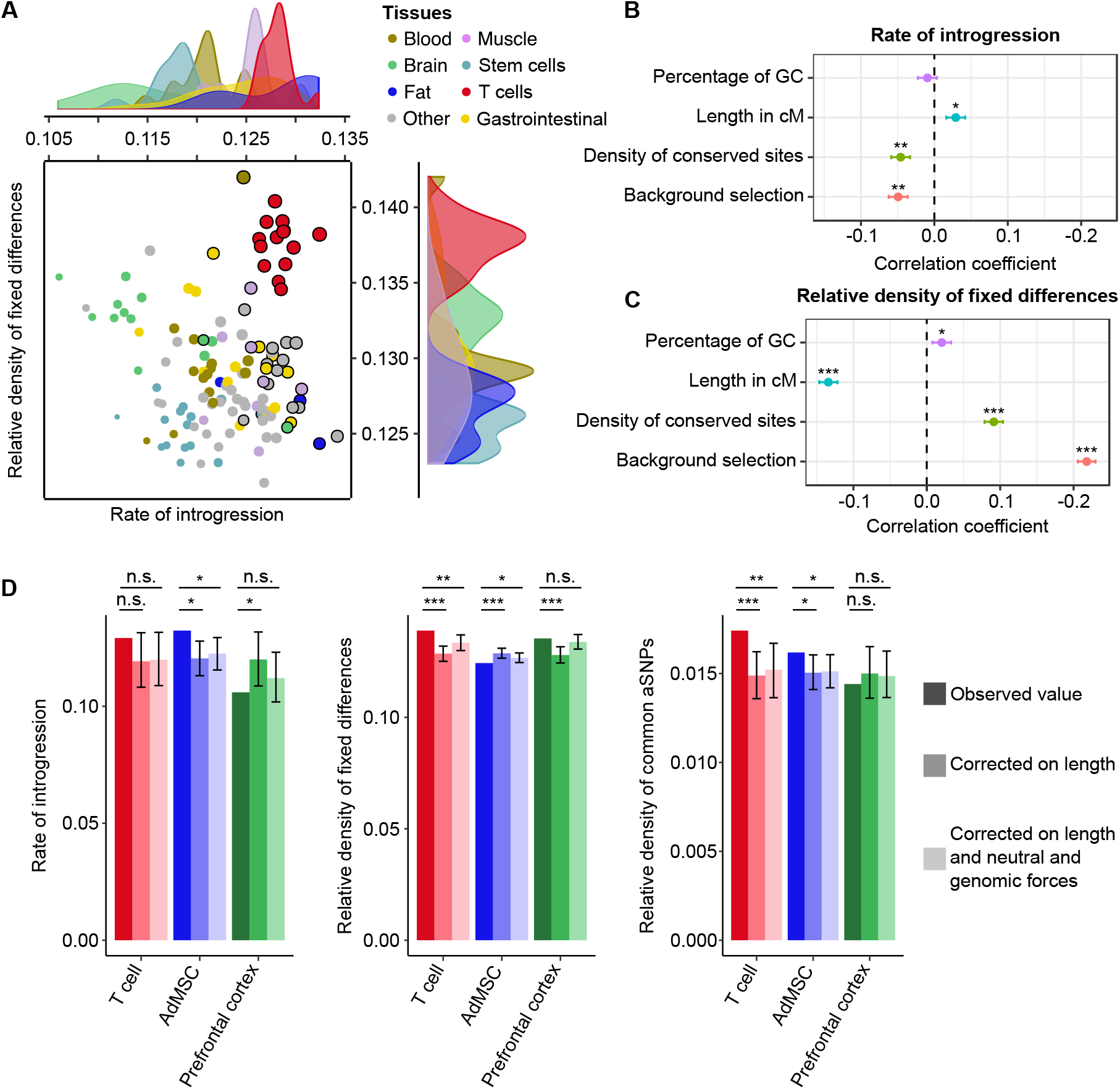
Factors shaping human-Neanderthal divergence and archaic introgression at enhancers. (A) Comparison of the relative density of human-Neanderthal divergence and rate of introgression in the enhancers of the 127 tissues studied. The size of the circles is proportional to the relative density of common aSNPs in the enhancers of the corresponding tissue; a black circle is added when the relative density of common aSNPs is significantly higher in these enhancers (FDR < 5%) than in the rest of the genome. The densities of each tissue category along the two axes are also presented. (B-C) Genome-wide correlations, using 100kb-windows, between the rate of Neanderthal introgression (B) or the relative density of fixed human-Neanderthal differences (C) and neutral and selective forces. * *p*-value < 10^-2^, ** *p*-value < 10^-10^, *** *p*-value < 10^-20^. (D) Observed values of relative density of fixed differences, introgression and common aSNPs observed at the enhancers of core T cells, AdMSC, and prefrontal cortex, with respect to expectations based on 100 kb windows matched for length of enhancers alone, or for length of enhancers, percentage of GC, recombination rate, density of conserved sites and mean B-statistic of their enhancers (see Supplemental Methods). n.s. not significant, * *p*-value < 0.05, ** *p*-value < 0.01, *** *p*-value < 10^-3^.

We then explored the factors that may drive, at the genome-wide level, the detected variation in Neanderthal divergence and archaic introgression. We correlated, using 100kb-windows, divergence and introgression with metrics that capture local variation in neutral (mutation, recombination) and selected (negative and background selection) diversity. Specifically, we measured the percentage of GC to account for their higher mutability, genetic size as measure of recombination rate, density of conserved sites (GerpRS > 2) as a measure of negative selection, and background selection derived as (1-B), where B is the mean B-statistic in the window.^34^ We found that background selection correlates with a lower rate of archaic introgression (r=-0.049, *p*-value <10^-15^, **Figure 4B**), consistently with previous findings,^1,2^ but also with increased local divergence (r = 0.22, *p*-value < 10^-20^, **Figure 4C**) and reduced density of both common variants and fixed differences (r = −0.46 and −0.05 respectively, *p*-value < 5×10^-20^, **Figure S1**). We also found that negative selection and recombination rate correlate with both divergence and introgression, even after adjusting for background selection (**Figure S2**).

To understand further how these factors could account for the variation in divergence and introgression detected at enhancers (**Figure S3**), we focused on 3 model tissues: T cells (enhancers with high divergence and introgression), AdMSC (enhancers with low divergence and high introgression), and prefrontal cortex (enhancers with high divergence and low introgression) (**Figure 4D**). When correcting for the various neutral and selective factors, introgression at T cell enhancers did not exceed that of other tissues (*p*-value > 0.11), but the high divergence and relative density of aSNPs remained significant (*p*-value < 8×10^-3^). For AdMSC, introgression remained higher than expected (*p*-value = 4×10^-3^), leading to an excess of aSNPs despite their depletion in divergence (*p*-value = 3.8×10^-2^). For enhancers active at prefrontal cortex, all variables were within expected bounds. Collectively, these analyses indicate that variation of several neutral and selective factors are not sufficient to explain the excess of Neanderthal introgression detected at enhancers. Some enhancers may have undergone past adaptation in the Neanderthal lineage or adaptive introgression in modern humans, as illustrated by T-cells and AdMSC, respectively.

Finally, we explored the impact of archaic introgression at enhancers on gene expression. To identify genes whose expression is altered by Neanderthal introgression at enhancers, we focused on tissues where promoter capture-HiC data were available^35,36^ and assigned each enhancer located in a promoter-interacting region to the corresponding gene(s). Archaic variants at enhancers predicted to interact with a gene were strongly enriched in eQTLs (OR=2.6, *p*-value<10^-3^, Supplemental Note 1 and **Figure S4**), supporting further the regulatory potential of aSNPs. Genes interacting with T cell enhancers that harbor common aSNPs (N = 1629, **Table S5**) were not enriched in any specific biological function. However, 285 of these genes are highly expressed in T cells (FPKM > 100), and include known regulators of the immune response (e.g., *CXCR4* [MIM: 162643], *IL7R* [MIM: 146661], *IL10RA* [MIM: 146933], *NFKBIA* [MIM: 164008] and *PTPRC* [MIM: 151460]). We found 14 loci presenting signatures of adaptive introgression; i.e., genes that interact with enhancers harboring very high frequency aSNPs (99^th^ percentile of MAF: MAF_CEU_ > 0.29 or MAF_CHB_ > 0.35; **Figures 5A,B**). Among these, we found *ANKRD27*, associated with eosinophilic esophagitis (MIM: 610247),^37^ and *MED15* (MIM: 607372), involved in several cancers.^38–40^ With respect to adipose-related tissues, we identified 690 genes — 43 of which being highly expressed (FPKM > 100) in the adipose tissue — interacting with AdMSC enhancers that contain common aSNPs (**Table S6**). These genes were enriched in functions related to the regulation of cell motility (GO:2000145, *p*-value < 2.0×10^-8^) and insulin-like growth factor binding protein complex (GO:0016942, *p*-value < 2.7×10^-5^) (**Table S7**). We detected 16 aSNPs at AdMSC enhancers that present strong signatures of adaptive introgression (**Figures 5C,D**).

**Figure 5.**
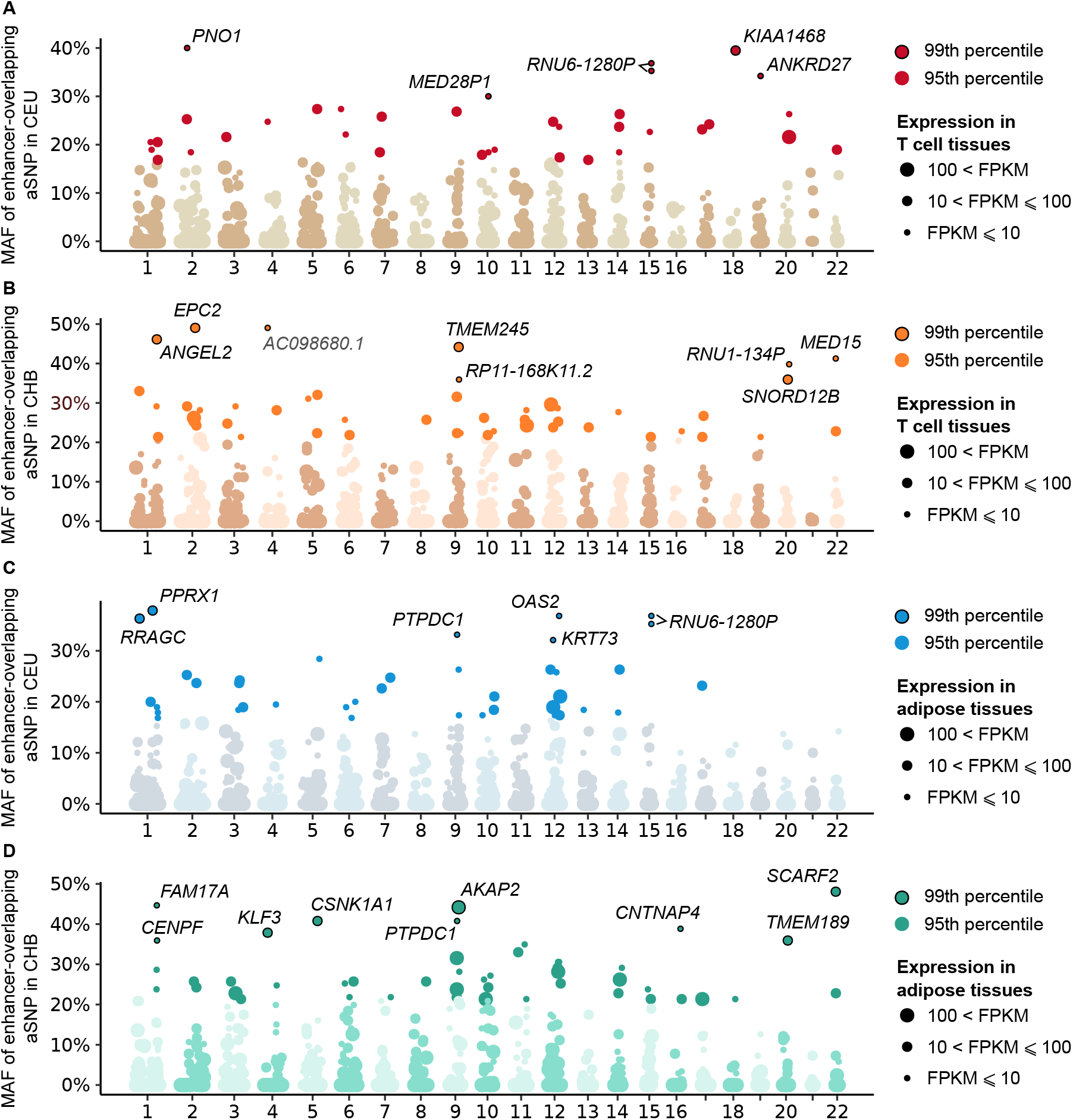
Manhattan plots of genes interacting with enhancers that contain archaic variants. (A-B) Genome-wide distribution of MAF in CEU or CHB at aSNPs that overlap enhancers active in T Cells (core T cell enhancers). For each window of 1Mb along the genome, only the aSNP with the highest MAF is shown. Point sizes reflect FPKM of the most expressed genes (max FPKM across T lymphocytes from Blueprint database^56^) among genes interacting with the enhancer in T cells.^35^ (C-D) Similar plots for enhancers active in AdMSC. Point sizes reflect the FPKM of the most expressed gene (max FPKM in GTEx tissues *Adipose – subcutaneous* and *Adipose – Visceral (Omentum)^57^)* among genes interacting with the enhancer in adipose tissue.^36^

## Discussion

This study reconstructs the history of how Neanderthal introgression has affected various types of regulatory elements as well as the mechanistic bases through which archaic variants have altered gene regulation. Although miRNAs *per se* harbor few archaic variants, some of them may impact strongly miRNA-mediated gene regulation and disease risk. For example, the archaic allele at miR-4293 (rs12220909) is responsible for the loss of 95% of its targets, and has been associated with diminished cancer susceptibility.^41,42^ Archaic introgression has also affected miRNA binding sites, as illustrated by *ONECUT2* (MIM: 604894), where an archaic haplotype that is present at high frequency in Asia (MAF_CHB_ = 0.49) alters multiple conserved miRNA binding sites. *ONECUT2* is involved in liver, pancreas and nervous system development,^43^ and has recently been proposed as a regulator of tumor growth in ovarian cancer (MIM: 167000).^44^

It is important to highlight that one third of the 127 tissues tested displayed a significant enrichment of archaic variants in their enhancers, reflecting either high human-Neanderthal differentiation, as observed in T-cell enhancers, or increased archaic introgression, as detected in AdMSC. Supporting the adaptive nature of the higher introgression detected at AdMSC, enhancers containing archaic variants interact preferentially with genes involved in the regulation of adipocyte differentiation and adipogenesis. These include receptors such as *PDGFRB* (MIM: 173410) and *TGFBR2* (MIM: 190182), the insulin growth factor *IGF1* (MIM: 147440) and its binding partners *IGFBP2* (MIM: 146731) and *IGFBP3* (MIM: 146732), or the *CXCR4* chemokine (MIM: 162643).^45–51^ Furthermore, two of the enhancers harboring archaic variants at the highest frequencies interact with key regulators of adipocyte differentiation, such as *KLF3* (MIM: 609392) and *PRRX1* (MIM: 167420).^52,53^ In view of the proposed adaptation of Neanderthals to cold environments,^54^ it is tempting to speculate that enhancers of adipose-derived mesenchymal stem cells carry today the remnants of such adaptive events. This hypothesis becomes particularly interesting in the light of previously reported cases of adaptive introgression at *LEPR* (MIM: 601007) and *WARS2/TBX15* (MIM: 604733/604127), both involved in the regulation of adipose tissue differentiation and body-fat distribution.^23,55^ Further studies aiming to functionally characterize the regulatory effects of Neanderthal variants on adipocyte differentiation and fat distribution are now warranted, as these archaic variants may have contributed to the adaptation of early Eurasians to colder climates.

## Supplemental Data

Supplemental Data include four figures, seven tables, supplemental Methods, and one Note.

## Acknowledgments

This work was supported by the *Institut Pasteur*, the *Centre Nationale de la Recherche Scientifique* (CNRS), and the *Agence Nationale de la Recherche* (ANR) grants: “IEIHSEER “ANR-14-CE14-0008-02 and “TBPATHGEN” ANR-14-CE14-0007-02. The laboratory of LQM has received funding from the French Government’s Investissement d’Avenir program, Laboratoire d’Excellence “Integrative Biology of Emerging Infectious Diseases” (grant no. ANR-10-LABX-62-IBEID).

## References

1. Sankararaman, S., Mallick, S., Dannemann, M., Prufer, K., Kelso, J., Paabo, S., Patterson, N., and Reich, D. (2014). The genomic landscape of Neanderthal ancestry in present-day humans. Nature 507, 354–357.

2. Sankararaman, S., Mallick, S., Patterson, N., and Reich, D. (2016). The Combined Landscape of Denisovan and Neanderthal Ancestry in Present-Day Humans. Curr Biol 26, 1241–1247.

3. Vernot, B., and Akey, J.M. (2014). Resurrecting surviving Neandertal lineages from modern human genomes. Science 343, 1017–1021.

4. Vernot, B., Tucci, S., Kelso, J., Schraiber, J.G., Wolf, A.B., Gittelman, R.M., Dannemann, M., Grote, S., McCoy, R.C., Norton, H., et al. (2016). Excavating Neandertal and Denisovan DNA from the genomes of Melanesian individuals. Science 352, 235–239.

5. Browning, S.R., Browning, B.L., Zhou, Y., Tucci, S., and Akey, J.M. (2018). Analysis of Human Sequence Data Reveals Two Pulses of Archaic Denisovan Admixture. Cell 173, 53–61.e59.

6. Prufer, K., Racimo, F., Patterson, N., Jay, F., Sankararaman, S., Sawyer, S., Heinze, A., Renaud, G., Sudmant, P.H., de Filippo, C., et al. (2014). The complete genome sequence of a Neanderthal from the Altai Mountains. Nature 505, 43–49.

7. Prufer, K., de Filippo, C., Grote, S., Mafessoni, F., Korlevic, P., Hajdinjak, M., Vernot, B., Skov, L., Hsieh, P., Peyregne, S., et al. (2017). A high-coverage Neandertal genome from Vindija Cave in Croatia. Science 358, 655–658.

8. Dannemann, M., and Kelso, J. (2017). The Contribution of Neanderthals to Phenotypic Variation in Modern Humans. Am J Hum Genet 101, 578–589.

9. Simonti, C.N., Vernot, B., Bastarache, L., Bottinger, E., Carrell, D.S., Chisholm, R.L., Crosslin, D.R., Hebbring, S.J., Jarvik, G.P., Kullo, I.J., et al. (2016). The phenotypic legacy of admixture between modern humans and Neandertals. Science 351, 737–741.

10. Dannemann, M., Prufer, K., and Kelso, J. (2017). Functional implications of Neandertal introgression in modern humans. Genome Biol 18, 61.

11. Quach, H., Rotival, M., Pothlichet, J., Loh, Y.E., Dannemann, M., Zidane, N., Laval, G., Patin, E., Harmant, C., Lopez, M., et al. (2016). Genetic Adaptation and Neandertal Admixture Shaped the Immune System of Human Populations. Cell 167, 643–656.

12. McCoy, R.C., Wakefield, J., and Akey, J.M. (2017). Impacts of Neanderthal-Introgressed Sequences on the Landscape of Human Gene Expression. Cell 168, 916–927.

13. Deschamps, M., Laval, G., Fagny, M., Itan, Y., Abel, L., Casanova, J.L., Patin, E., and Quintana-Murci, L. (2016). Genomic Signatures of Selective Pressures and Introgression from Archaic Hominins at Human Innate Immunity Genes. Am J Hum Genet 98, 5–21.

14. Enard, D., and Petrov, D.A. (2018). Evidence that RNA Viruses Drove Adaptive Introgression between Neanderthals and Modern Humans. Cell 175, 360–371.

15. Sams, A.J., Dumaine, A., Nedelec, Y., Yotova, V., Alfieri, C., Tanner, J.E., Messer, P.W., and Barreiro, L.B. (2016). Adaptively introgressed Neandertal haplotype at the OAS locus functionally impacts innate immune responses in humans. Genome Biol 17, 246.

16. Jegou, B., Sankararaman, S., Rolland, A.D., Reich, D., and Chalmel, F. (2017). Meiotic Genes Are Enriched in Regions of Reduced Archaic Ancestry. Mol Biol Evol 34, 1974–1980.

17. Kuhlwilm, M., Gronau, I., Hubisz, M.J., de Filippo, C., Prado-Martinez, J., Kircher, M., Fu, Q., Burbano, H.A., Lalueza-Fox, C., de la Rasilla, M., et al. (2016). Ancient gene flow from early modern humans into Eastern Neanderthals. Nature 530, 429–433.

18. Rogers, A.R., Bohlender, R.J., and Huff, C.D. (2017). Early history of Neanderthals and Denisovans. Proc Natl Acad Sci U S A 114, 9859–9863.

19. Harris, K., and Nielsen, R. (2016). The Genetic Cost of Neanderthal Introgression. Genetics 203, 881–891.

20. Juric, I., Aeschbacher, S., and Coop, G. (2016). The Strength of Selection against Neanderthal Introgression. PLoS Genet 12, e1006340.

21. Petr, M., Pääbo, S., Kelso, J., and Vernot, B. (2018). The limits of long-term selection against Neandertal introgression. bioRxiv doi.org/10.1101/362566

22. Racimo, F., Marnetto, D., and Huerta-Sanchez, E. (2017). Signatures of Archaic Adaptive Introgression in Present-Day Human Populations. Mol Biol Evol 34, 296–317.

23. Racimo, F., Gokhman, D., Fumagalli, M., Ko, A., Hansen, T., Moltke, I., Albrechtsen, A., Carmel, L., Huerta-Sanchez, E., and Nielsen, R. (2017). Archaic Adaptive Introgression in TBX15/WARS2. Mol Biol Evol 34, 509–524.

24. Gittelman, R.M., Schraiber, J.G., Vernot, B., Mikacenic, C., Wurfel, M.M., and Akey, J.M. (2016). Archaic Hominin Admixture Facilitated Adaptation to Out-of-Africa Environments. Curr Biol 26, 3375–3382.

25. Racimo, F., Sankararaman, S., Nielsen, R., and Huerta-Sanchez, E. (2015). Evidence for archaic adaptive introgression in humans. Nat Rev Genet 16, 359–371.

26. Huerta-Sanchez, E., Jin, X., Asan, Bianba, Z., Peter, B.M., Vinckenbosch, N., Liang, Y., Yi, X., He, M., Somel, M., et al. (2014). Altitude adaptation in Tibetans caused by introgression of Denisovan-like DNA. Nature 512, 194–197.

27. Jagoda, E., Lawson, D.J., Wall, J.D., Lambert, D., Muller, C., Westaway, M., Leavesley, M., Capellini, T.D., Mirazon Lahr, M., Gerbault, P., et al. (2017). Disentangling Immediate Adaptive Introgression from Selection on Standing Introgressed Variation in Humans. Mol Biol Evol 35, 623–630.

28. Dannemann, M., and Racimo, F. (2018). Something old, something borrowed: admixture and adaptation in human evolution. Curr Opin Genet Dev 53, 1–8.

29. Kudaravalli, S., Veyrieras, J.B., Stranger, B.E., Dermitzakis, E.T., and Pritchard, J.K. (2009). Gene expression levels are a target of recent natural selection in the human genome. Mol Biol Evol 26, 649–658.

30. Villar, D., Berthelot, C., Aldridge, S., Rayner, T.F., Lukk, M., Pignatelli, M., Park, T.J., Deaville, R., Erichsen, J.T., Jasinska, A.J., et al. (2015). Enhancer evolution across 20 mammalian species. Cell 160, 554–566.

31. Roadmap Epigenomics Consortium, Kundaje, A., Meuleman, W., Ernst, J., Bilenky, M., Yen, A., Heravi-Moussavi, A., Kheradpour, P., Zhang, Z., Wang, J., et al. (2015). Integrative analysis of 111 reference human epigenomes. Nature 518, 317–330.

32. Enright, A.J., John, B., Gaul, U., Tuschl, T., Sander, C., and Marks, D.S. (2003). MicroRNA targets in Drosophila. Genome Biol 5, R1.

33. 1000 Genomes Project Consortium, Auton, A., Brooks, L.D., Durbin, R.M., Garrison, E.P., Kang, H.M., Korbel, J.O., Marchini, J.L., McCarthy, S., McVean, G.A., et al. (2015). A global reference for human genetic variation. Nature 526, 68–74.

34. McVicker, G., Gordon, D., Davis, C., and Green, P. (2009). Widespread genomic signatures of natural selection in hominid evolution. PLoS Genet 5, e1000471.

35. Javierre, B.M., Burren, O.S., Wilder, S.P., Kreuzhuber, R., Hill, S.M., Sewitz, S., Cairns, J., Wingett, S.W., Varnai, C., Thiecke, M.J., et al. (2016). Lineage-Specific Genome Architecture Links Enhancers and Non-coding Disease Variants to Target Gene Promoters. Cell 167, 1369–1384.

36. Pan, D.Z., Garske, K.M., Alvarez, M., Bhagat, Y.V., Boocock, J., Nikkola, E., Miao, Z., Raulerson, C.K., Cantor, R.M., Civelek, M., et al. (2018). Integration of human adipocyte chromosomal interactions with adipose gene expression prioritizes obesity-related genes from GWAS. Nat Commun 9, 1512.

37. Sleiman, P.M., Wang, M.L., Cianferoni, A., Aceves, S., Gonsalves, N., Nadeau, K., Bredenoord, A.J., Furuta, G.T., Spergel, J.M., and Hakonarson, H. (2014). GWAS identifies four novel eosinophilic esophagitis loci. Nat Commun 5, 5593.

38. Syring, I., Weiten, R., Muller, T., Schmidt, D., Steiner, S., Kristiansen, G., Muller, S.C., and Ellinger, J. (2018). The knockdown of the Mediator complex subunit MED15 restrains urothelial bladder cancer cells’ malignancy. Oncol Lett 16, 3013–3021.

39. Weiten, R., Muller, T., Schmidt, D., Steiner, S., Kristiansen, G., Muller, S.C., Ellinger, J., and Syring, I. (2018). The Mediator complex subunit MED15, a promoter of tumour progression and metastatic spread in renal cell carcinoma. Cancer Biomark 21, 839–847.

40. Shaikhibrahim, Z., Offermann, A., Halbach, R., Vogel, W., Braun, M., Kristiansen, G., Bootz, F., Wenzel, J., Mikut, R., Lengerke, C., et al. (2015). Clinical and molecular implications of MED15 in head and neck squamous cell carcinoma. Am J Pathol 185, 1114–1122.

41. Fan, L., Chen, L., Ni, X., Guo, S., Zhou, Y., Wang, C., Zheng, Y., Shen, F., Kolluri, V.K., Muktiali, M., et al. (2017). Genetic variant of miR-4293 rs12220909 is associated with susceptibility to non-small cell lung cancer in a Chinese Han population. PLoS One 12, e0175666.

42. Zhang, P., Wang, J., Lu, T., Wang, X., Zheng, Y., Guo, S., Yang, Y., Wang, M., Kolluri, V.K., Qiu, L., et al. (2015). miR-449b rs10061133 and miR-4293 rs12220909 polymorphisms are associated with decreased esophageal squamous cell carcinoma in a Chinese population. Tumour Biol 36, 8789–8795.

43. Kropp, P.A., and Gannon, M. (2016). Onecut transcription factors in development and disease. Trends Dev Biol 9, 43–57.

44. Lu, T., Wu, B., Yu, Y., Zhu, W., Zhang, S., Zhang, Y., Guo, J., and Deng, N. (2018). Blockade of ONECUT2 expression in ovarian cancer inhibited tumor cell proliferation, migration, invasion and angiogenesis. Cancer Sci 109, 2221–2234.

45. Gao, Z., Daquinag, A.C., Su, F., Snyder, B., and Kolonin, M.G. (2018). PDGFRalpha/PDGFRbeta signaling balance modulates progenitor cell differentiation into white and beige adipocytes. Development 145, dev155861.

46. Onogi, Y., Wada, T., Kamiya, C., Inata, K., Matsuzawa, T., Inaba, Y., Kimura, K., Inoue, H., Yamamoto, S., Ishii, Y., et al. (2017). PDGFRbeta Regulates Adipose Tissue Expansion and Glucose Metabolism via Vascular Remodeling in Diet-Induced Obesity. Diabetes 66, 1008–1021.

47. Kim, Y.J., Hwang, S.J., Bae, Y.C., and Jung, J.S. (2009). MiR-21 regulates adipogenic differentiation through the modulation of TGF-beta signaling in mesenchymal stem cells derived from human adipose tissue. Stem Cells 27, 3093–3102.

48. Chang, H.R., Kim, H.J., Xu, X., and Ferrante, A.W., Jr. (2016). Macrophage and adipocyte IGF1 maintain adipose tissue homeostasis during metabolic stresses. Obesity (Silver Spring) 24, 172–183.

49. Yau, S.W., Russo, V.C., Clarke, I.J., Dunshea, F.R., Werther, G.A., and Sabin, M.A. (2015). IGFBP-2 inhibits adipogenesis and lipogenesis in human visceral, but not subcutaneous, adipocytes. Int J Obes (Lond) 39, 770–781.

50. Chan, S.S., Schedlich, L.J., Twigg, S.M., and Baxter, R.C. (2009). Inhibition of adipocyte differentiation by insulin-like growth factor-binding protein-3. Am J Physiol Endocrinol Metab 296, E654–663.

51. Yao, L., Heuser-Baker, J., Herlea-Pana, O., Zhang, N., Szweda, L.I., Griffin, T.M., and Barlic-Dicen, J. (2014). Deficiency in adipocyte chemokine receptor CXCR4 exacerbates obesity and compromises thermoregulatory responses of brown adipose tissue in a mouse model of diet-induced obesity. FASEB J 28, 4534–4550.

52. Sue, N., Jack, B.H., Eaton, S.A., Pearson, R.C., Funnell, A.P., Turner, J., Czolij, R., Denyer, G., Bao, S., Molero-Navajas, J.C., et al. (2008). Targeted disruption of the basic Kruppel-like factor gene (Klf3) reveals a role in adipogenesis. Mol Cell Biol 28, 3967–3978.

53. Du, B., Cawthorn, W.P., Su, A., Doucette, C.R., Yao, Y., Hemati, N., Kampert, S., McCoin, C., Broome, D.T., Rosen, C.J., et al. (2013). The transcription factor paired-related homeobox 1 (Prrx1) inhibits adipogenesis by activating transforming growth factor-beta (TGFbeta) signaling. J Biol Chem 288, 3036–3047.

54. Steegmann, A.T., Jr., Cerny, F.J., and Holliday, T.W. (2002). Neandertal cold adaptation: physiological and energetic factors. Am J Hum Biol 14, 566–583.

55. Sazzini, M., Schiavo, G., De Fanti, S., Martelli, P.L., Casadio, R., and Luiselli, D. (2014). Searching for signatures of cold adaptations in modern and archaic humans: hints from the brown adipose tissue genes. Heredity (Edinb) 113, 259–267.

56. Chen, L., Ge, B., Casale, F.P., Vasquez, L., Kwan, T., Garrido-Martin, D., Watt, S., Yan, Y., Kundu, K., Ecker, S., et al. (2016). Genetic Drivers of Epigenetic and Transcriptional Variation in Human Immune Cells. Cell 167, 1398–1414 e1324.

57. GTEx Consortium (2013). The Genotype-Tissue Expression (GTEx) project. Nat Genet 45, 580–585.

