## Supplemental files for "Impact and evolutionary determinants of Neanderthal introgression on transcriptional and post-transcriptional regulation"

### Supplemental Figures

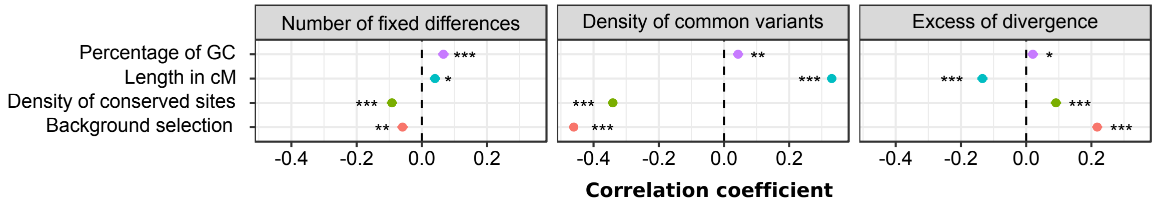


**Figure S1. Effects of neutral and selective factors on the density of common variants and fixed human-Neanderthal differences.** Genome-wide correlations, using 100kb-windows, between the density of fixed human-Neanderthal differences, the density of common variants in Eurasia or the ratio of these quantities and several proxies of neutral and selective factors. *$p$-value < $10^{-2}$, **$p$-value < $10^{-10}$, ***$p$-value < $10^{-20}$.

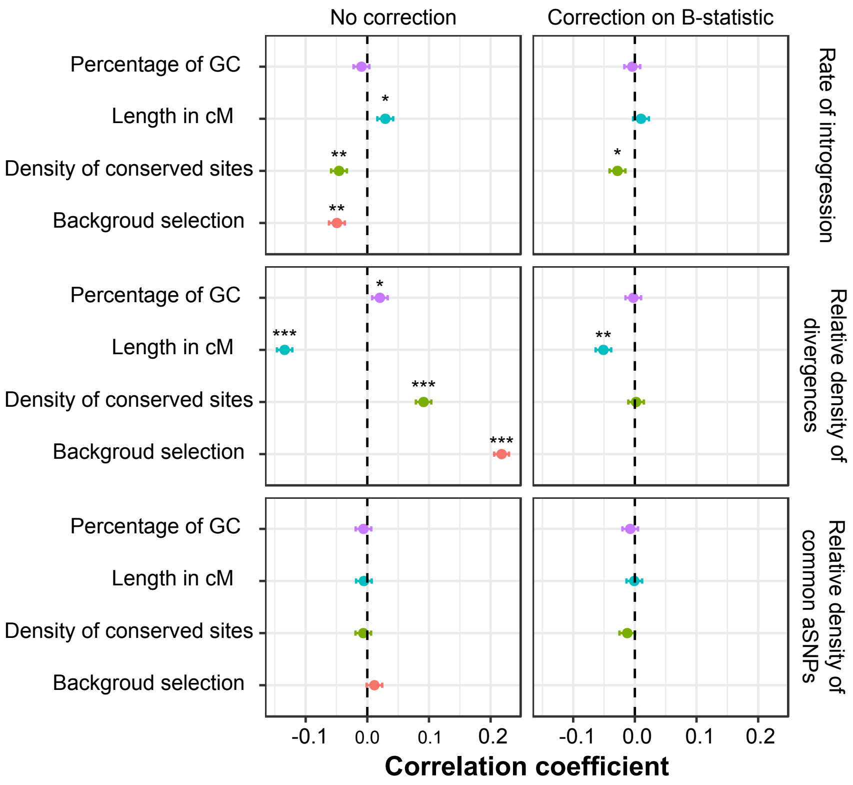


**Figure S2. Effects of neutral and selective factors on rate of introgression and relative density of fixed human-Neanderthal differences, conditional on background selection.** Correlations, computed in 100kb windows along the genome, between the rate of introgression, the relative density of fixed human-Neanderthal differences and common aSNPs, and several proxies of neutral and selective factors. *$p$-value < $10^{-2}$, **$p$-value < $10^{-10}$, ***$p$-value < $10^{-20}$.

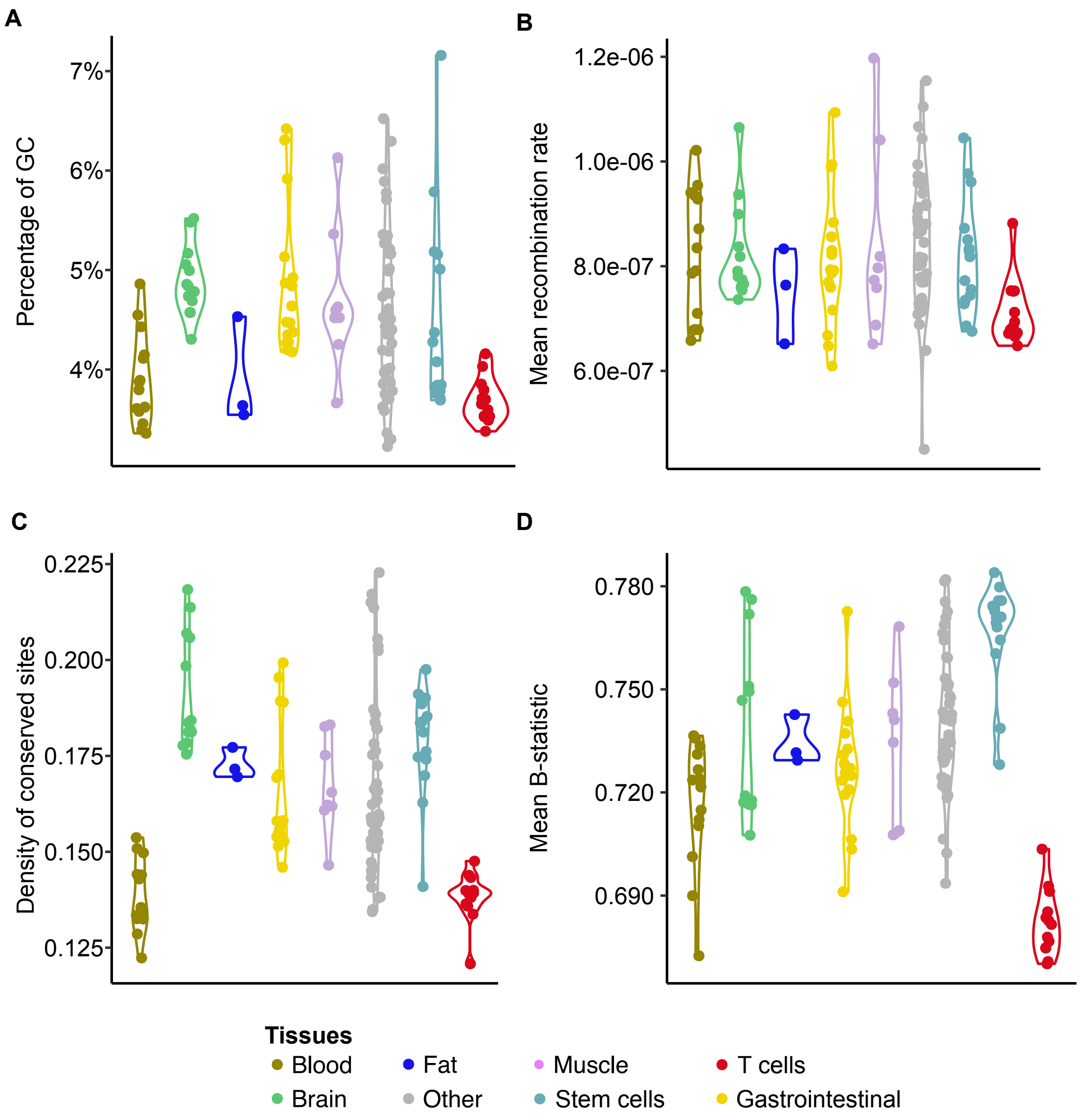


**Figure S3. Intensity of neutral and selective factors in enhancers across tissues.** Values, in the enhancers of the 127 tissues studied, of the percentage of GC, the mean recombination rate, the density of conserved sites (GerpRS > 2), and the mean B-statistic.

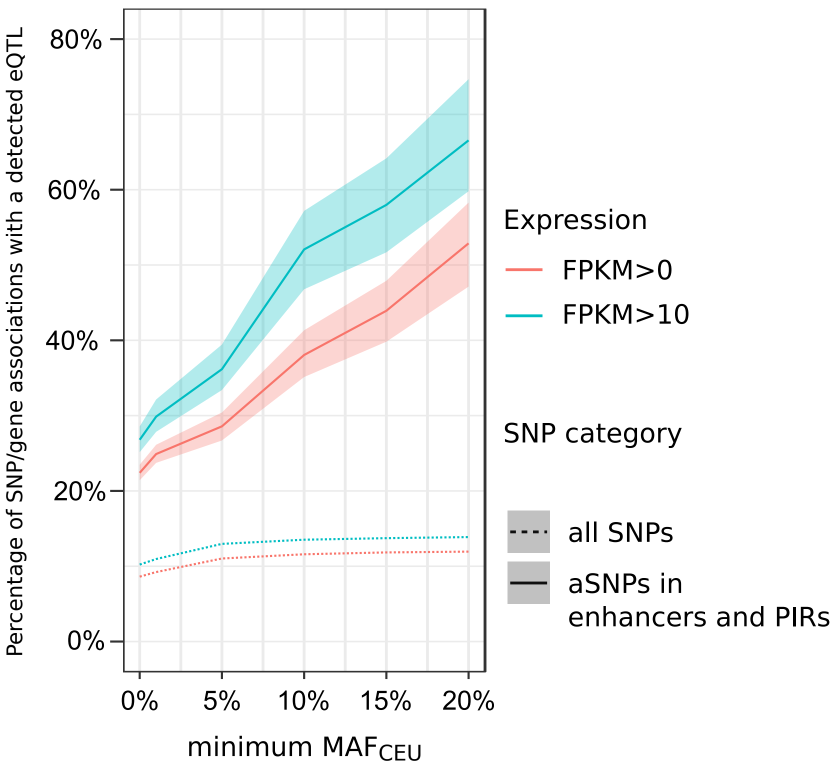


**Figure S4. Effect of enhancer variants on gene expression.** Comparison of the proportion of SNP/gene pairs with an eQTL in the eQTLGen Consortium,[1] as a function of the frequency of the SNP in the CEU population, and the expression of the gene in whole blood[2] for two classes of SNPs: all SNPs tested in eQTLGen dataset (dotted line), and aSNPs that are in T cell enhancers that interact with the gene promoter (i.e. promoter interacting region – PIR) based on T cell contact maps[3] (plain line). Shaded regions indicate 95% confidence intervals computed by bootstrap.

### Supplemental Methods

#### Definition of archaic SNPs (aSNPs)

We considered all SNPs present in the European (CEU) and Asian (CHB) populations of the 1000 Genomes Consortium phase 3 (ref.[4]). Among them, aSNPs were defined as SNPs that (i) have an allele for which the Neanderthal Altai is homozygous,[5] (ii) are absent from the African Yoruba population, and (iii) are located in a region in which Neanderthal introgression has already been detected in Eurasia (probability of Neanderthal introgression > 0.9) (ref.[6]). Because variants due to incomplete lineage sorting are more likely to segregate at high frequency, and to minimize false positives presenting signals of adaptive introgression, we took additional steps to filter out such variants when considering aSNPs at high frequencies (**Figure 5** and **Tables S5** and **S6**). Specifically, we retrieved for each aSNP the set of all aSNPs that are in high linkage disequilibrium ($r^2 > 0.8$) in either CEU or CHB. We then required variants to have at least one linked aSNP at a distance of >10 kb, thus filtering likely cases of incomplete lineage sorting,

#### Relative density of aSNPs and enrichments

To measure the impact of Neanderthal introgression on a specific region, or set of regions, we measured the density of Neanderthal variants, as the number of aSNPs in the region, divided by the length (in bp) of the study region. Likewise, the density of non-archaic variants was computed as a measure of the overall diversity of the region. We then measured the excess or depletion of archaic variants in a region by computing the ratios of these densities (i.e. relative density of aSNPs) in the region, which were compared with those of the rest of the genome. In doing so, we obtained an odds ratio that is significantly higher than 1 if the region presents an excess of aSNPs, and significantly lower than 1 if the region is depleted in aSNPs. We also used a variation of this statistic considering only aSNPs and SNPs within a given range of frequencies *f (*MAF<1%, 1% ≤ MAF ≤ 5%, or MAF > 5**%**).

$$\frac{\#aSNPs^{f}_{In\ Region} \Big/ \#classical\ SNPs^{f}_{in\ Region}}{\#aSNPs^{f}_{Outside\ Region} \Big/ \#classical\ SNPs^{f}_{Oustide\ Region}}$$

To compute the significance of the odds ratio, while considering both the haplotype structure of Neanderthal variants and the local structure of the study regions, we divided the genome into windows of 100kb and performed 10,000 bootstrap resamples of these windows, recomputing the odds ratio for each bootstrap sample. We then computed enrichment/depletion *p*-values as the percentage of bootstrap resamples where the odds ratio is lower/higher than 1. Bidirectional *p*-values were then obtained as 2×min($p_{enrichment}$, $p_{depletion}$)

**Definition of regulatory regions**

Human miRNA sequences and their locations were obtained from the miRbase database, version 20 (ref.[7]). We used the miRanda software[8] version 3.3a, to predict miRNA binding sites in the 3'UTR of coding genes, as defined in Ensembl Annotation GRcH37.70. Defaults cutoffs were used. Promoters and enhancers were defined based on chromatin marks in the 127 tissues of the Roadmap Epigenomics Consortium.[9] The calling of promoters and enhancers was performed based on 15-state ChromHMM.[10] We considered the union of the *Active TSS* and *Flanking* TSS as "promoters", and the union of the *Enh* (enhancers) and *EnhG* (enhancers genic) categories as "enhancers".

**Characterizing the impact of introgression on regulatory regions**

To dissect the relative contribution of Human-Neanderthal divergence, and post-admixture removal of Neanderthal introgressed variants in shaping the current landscape of introgressed regulatory variants, we first searched for fixed differences between the genomes of Neanderthals and modern humans. Namely, we considered as a fixed difference any variant (i) where both Neanderthal Altai[5] and Neanderthal Vindija[11] were homozygous for an allele, (ii) absent in 6EPO alignments and (iii) absent in the Yoruba population.[4] We then defined the density of fixed differences in a region as the number of fixed differences over the number of sites in that region, where sequence information was available for Altai, Vindija and 6EPO genomes. This density was further divided by the density of common variants in the region to yield a 'relative density of fixed differences', which measures the excess of divergence in a study region given its overall diversity. Reciprocally, we considered as the rate of introgression, the percentage of fixed differences that were introgressed into modern humans and reach a MAF of at least 5%. With these definitions, the product of the rate of introgression and the relative density of fixed differences is equal to the relative density of common aSNPs in the region.

**Impact of neutral and selective factors on introgression-related metrics**

We investigated the effects of mutation, recombination, and negative selection (directly or indirectly through background selection) on various introgression-related metrics, including the rate of introgression, the relative density of fixed differences, as well as the density of archaic variants segregating in CEU and CHB populations. To do so, we split the human genome into 100kb windows, and focused on sites where sequence information was available for Altai, Vindija and 6EPO genomes, excluding windows where sequence information was available for less than 50% of the windows. We then computed, for each window, the percentage of GC or CG dinucleotide in the sequence, the mean recombination rate, the proportion of conserved sites (GerpRS > 2) and the mean B-statistic. For each of these metrics, the Pearson's correlations with each introgression-related metric were computed across all windows.

Next, similarly to what we performed genome-wide, we subdivided the genome in 100kb windows and, for each tissue, we computed, at windows containing enhancers, the total enhancer's length and the percentage of GC, mean recombination rate, percentage of conserved sites (GerpRS >2) and mean B-statistic in the corresponding enhancers. For each tissue, we then assembled enhancers from randomly sampled windows and tissues to create a pseudo-tissue, for which we can compute the relative density of fixed differences, the rate of introgression and relative density of common aSNPs. To ensure that the reconstructed tissues had an enhancer structure that is comparable to the original tissue, each resampled window and tissue were selected so that the length of their enhancers matched that of the enhancers from the original tissue.

To evaluate the contribution of neutral and selective forces to the relative density of fixed differences and rate of introgression, we performed additional resamples matching enhancers simultaneously for their percentage of GC, mean recombination rate, percentage of conserved sites and mean B-statistic, in addition to their length. For each tested tissue and matching, a total of 1000 resampling was performed and a *p*-value was computed as the number of resamplings for which the relative density of fixed differences or rate of introgression at enhancers of the tested tissue exceeded that of enhancers in the reconstructed tissue. When resampling, we used the following bins for matching: (i) total enhancer length: 20 bins defined as follows [0-200 bp], ]200-400 bp], ]400-600 bp], ]600-800 bp], ]800 bp-1kb], ]1-1.5 kb], ]1.5-2 kb], ]2-3 kb], ]3-4 kb], ]4-5 kb], ]5-7.5 kb], ]7.5-10 kb], ]10-20 kb], ]20-30 kb], ]30-40 kb], ]40-50 kb], ]50-75 kb], ]75-100 kb], and ]100-200 kb], (ii) percentage of GC and percentage of sites with GerpRS > 2: 20 uniformly distributed bins of 5% width, (iii) B-statistic: 10 uniform bins of width 0.1, and (iv) mean recombination rate: 10 bins, based on deciles.

### Identification of enhancer-interacting genes

To assign genes to the enhancers detected that are active in AdMSC, we used promoter-capture HiC (PC-HiC) data obtained from adipose tissue,[12] and assigned each promoter to a gene when it is located within 100 bp of its TSS. We then selected all interactions with a CHiCAGO score above 5, where the promoter-interacting region overlapped an enhancer in

AdMSC, and assigned the corresponding genes as enhancer's targets. For primary T cells, we used PC-HiC data obtained from Javierre *et al*.[3] We selected interactions with CHiCAGO score above 5 in the total $CD8^+$ T cells, as promoter interacting regions in this cell type showed the strongest overlap with core T cell enhancers (Jaccard Index = 9.7%).

**GO Enrichments**

To assess whether specific biological functions had been preferentially affected by archaic introgression at enhancers, we considered both tissues where PC-HiC was available, and assigned each enhancer to a gene based on promoter interactions. As enhancers can control multiple genes (22% of core T Cell enhancers are associated to more that 5 genes, with up to 73 associated genes for the same enhancer), and genes that share a common biological function tend to be found in clusters along the genome, we filtered out enhancers with more than 3 target genes from our enrichment analysis, thus reducing the risk of spurious enrichments due to clusters of co-regulated genes. We then used the GOseq package[13] to search for biological functions overrepresented among genes with aSNPs in their enhancers, using the set of all genes with a SNP in their enhancers as background and adjusting on total enhancer length of each gene.

### Supplemental Note 1: Effect of aSNPs in enhancers on gene expression

To assess the impact on gene expression of aSNPs that overlap enhancer regions, we first considered, for each gene, the set of aSNPs that overlap promoter-interacting enhancers in primary T-cells (focusing on core T cell enhancers). We then assessed the frequency at which such aSNPs were associated with changes in gene expression, based on GTEx eQTLs and whole blood eQTLs identified by the eQTLGen consortium.[1,2] We found that while only ~1% of aSNPs that overlap core T cell enhancers regulate their associated gene in GTEx tissues (FDR <5%), this figure reaches 22% when considering eQTLs obtained through metanalysis of whole blood samples from over 30,000 donors.[1] This suggests that while enhancer-overlapping aSNPs contribute to gene expression variability, large sample sizes are required to assess their true effects. Consistent with this notion, we observed that the proportion of enhancer aSNPs that control the expression of their associated genes increases with median gene expression and allele frequency (**Figure S4**), reaching 67% for genes with FPKM>10 and aSNPs with a MAF >20% in Europe. Our data suggests that while >60% of enhancer-overlapping aSNPs are significantly associated with gene expression variation, many of these associations are usually missed by eQTL studies due to low power or under-representation of individuals of non-European ancestry.

### Supplemental References


1. Võsa, U., Claringbould, A., Westra, H.-J., Bonder, M.J., Deelen, P., Zeng, B., Kirsten, H., Saha, A., Kreuzhuber, R., Kasela, S., et al. (2018). Unraveling the polygenic architecture of complex traits using blood eQTL meta-analysis. bioRxiv, doi.org/10.1101/447367
2. GTEx Consortium (2013). The Genotype-Tissue Expression (GTEx) project. Nat Genet *45*, 580-585.
3. Javierre, B.M., Burren, O.S., Wilder, S.P., Kreuzhuber, R., Hill, S.M., Sewitz, S., Cairns, J., Wingett, S.W., Varnai, C., Thiecke, M.J., et al. (2016). Lineage-Specific Genome Architecture Links Enhancers and Non-coding Disease Variants to Target Gene Promoters. Cell *167*, 1369-1384.
4. 1000 Genomes Project Consortium, Auton, A., Brooks, L.D., Durbin, R.M., Garrison, E.P., Kang, H.M., Korbel, J.O., Marchini, J.L., McCarthy, S., McVean, G.A., et al. (2015). A global reference for human genetic variation. Nature *526*, 68-74.
5. Prufer, K., Racimo, F., Patterson, N., Jay, F., Sankararaman, S., Sawyer, S., Heinze, A., Renaud, G., Sudmant, P.H., de Filippo, C., et al. (2014). The complete genome sequence of a Neanderthal from the Altai Mountains. Nature *505*, 43-49.
6. Sankararaman, S., Mallick, S., Dannemann, M., Prufer, K., Kelso, J., Paabo, S., Patterson, N., and Reich, D. (2014). The genomic landscape of Neanderthal ancestry in present-day humans. Nature *507*, 354-357.
7. Chou, C.H., Shrestha, S., Yang, C.D., Chang, N.W., Lin, Y.L., Liao, K.W., Huang, W.C., Sun, T.H., Tu, S.J., Lee, W.H., et al. (2018). miRTarBase update 2018: a resource for experimentally validated microRNA-target interactions. Nucleic Acids Res *46*, D296-D302.
8. Enright, A.J., John, B., Gaul, U., Tuschl, T., Sander, C., and Marks, D.S. (2003). MicroRNA targets in Drosophila. Genome Biol *5*, R1.
9. Roadmap Epigenomics Consortium, Kundaje, A., Meuleman, W., Ernst, J., Bilenky, M., Yen, A., Heravi-Moussavi, A., Kheradpour, P., Zhang, Z., Wang, J., et al. (2015). Integrative analysis of 111 reference human epigenomes. Nature *518*, 317-330.
10. Ernst, J., and Kellis, M. (2017). Chromatin-state discovery and genome annotation with ChromHMM. Nat Protoc *12*, 2478-2492.
11. Prufer, K., de Filippo, C., Grote, S., Mafessoni, F., Korlevic, P., Hajdinjak, M., Vernot, B., Skov, L., Hsieh, P., Peyregne, S., et al. (2017). A high-coverage Neandertal genome from Vindija Cave in Croatia. Science *358*, 655-658.
12. Pan, D.Z., Garske, K.M., Alvarez, M., Bhagat, Y.V., Boocock, J., Nikkola, E., Miao, Z., Raulerson, C.K., Cantor, R.M., Civelek, M., et al. (2018). Integration of human adipocyte chromosomal interactions with adipose gene expression prioritizes obesity-related genes from GWAS. Nat Commun *9*, 1512.
13. Young, M.D., Wakefield, M.J., Smyth, G.K., and Oshlack, A. (2010). Gene ontology analysis for RNA-seq: accounting for selection bias. Genome Biol *11*, R14.
